# Early steps of multi-receptor viral interactions dissected by high-density, multi-color single molecule mapping in living cells

**DOI:** 10.1101/2023.09.29.560105

**Authors:** Nicolas Mateos, Enric Gutierrez-Martinez, Jessica Angulo-Capel, Irene Carlon-Andres, Sergi Padilla-Parra, Maria F. Garcia-Parajo, Juan Andres Torreno-Pina

**Author notes:** Corresponding authors: Maria F. Garcia-Parajo, Juan Andres Torreno-Pina.

## Abstract

Direct visualization of the early steps of multi-receptor viral interactions at the singlemolecule level has been largely impeded by the technical challenges associated to imaging individual multi-molecular systems at relevant spatial (nanometer) and temporal (millisecond) scales. Here, we present a four-color, high-density single-molecule spatiotemporal mapping methodology to capture real-time interactions between individual viruses and three different viral (co-)receptors on the membrane of living immune cells derived from donors. Together with quantitative tools, our approach revealed the existence of a coordinated spatiotemporal diffusion of the three different (co-)receptors prior to viral-engagement. By varying the temporal-windows of cumulated single-molecule localizations, we discovered that such a concerted diffusion impacts on the residence time of viruses on the host membrane and potential viral infectivity. Overall, our methodology opens a new door to the systematic analysis of the initial steps of viral-host interactions and paves the way to the investigation of other multi-molecular systems at the single-molecule level.

## Introduction

Virus-mediated infectious diseases evolving into pandemics pose a recurrent challenge to public health worldwide. Prominent examples are the HIV-1 and SARS-CoV-2 viruses that have resulted in AIDS and COVID-19 pandemics, respectively. As such, huge efforts are being deployed in the development of vaccines and other therapeutic strategies against these, and other viral infections^1,2,3,4^. Along these lines, understanding the early molecular events leading to viral capture and cell entry is probably the most crucial step towards designing and developing vaccines able to elicit neutralizing antibodies against the different viral strains. Hence, the development of novel techniques able to capture the early events of viral infections is of urgent need.

Single virus tracking (SVT) has been widely and successfully used in the last two decades to study viral infections with unprecedented levels of details at the relevant spatiotemporal scales^5,6,7,8^. Indeed, by following the fate of individual viruses, SVT has shed light into how viruses interact and infect host cells by relying on a variety of cellular processes such as membrane-fusion^9^, clathrin or caveolin-mediated endocytosis^10, 11, 12, 13, 14, 15^ and cytoskeleton-dependent intracellular transport^10, 16, 17, 18, 19^. Nevertheless and despite these advances, still little is known about the first initial interactions between the virus and target membrane receptors since studying multi-molecular systems at the single molecule level with nanometer-spatial and millisecond-temporal resolutions remains highly challenging.

Dendritic cells (DCs) are antigen presenting cells of the immune system whose mission is to capture antigens in peripheral tissues and present them to T cells in lymphoid tissues^20, 21, 22^. DCs recognize and capture antigens by expressing a large number of pathogen recognition receptors on their cell membrane. A prominent example is DC-SIGN, a C-type lectin expressed on immature DCs (imDCs) with a carbon-recognition domain (CRD) able to bind and capture a large plethora of pathogens, including HIV-1^23^. Although the major targets of HIV-1 infection are CD4^+^-T cells where infection occurs via membrane-fusion^24^, it has been extensively shown that DCs crucially contribute to HIV-1 infection by influencing viral transmission and CD4^+^-T cell target infection^23, 25^. This process is initiated by the capture of HIV-1 by DC-SIGN on imDCs and subsequent DC-SIGN-mediated endocytosis by clathrin-dependent and -independent pathways^26, 27^. Importantly, it has been recently proposed that DC-SIGN also mediates the capture of SARS-CoV-2 viruses although the entry mechanisms are yet to be elucidated^28^. By applying single particle tracking (SPT) and super-resolution microscopy, we recently showed that the spatiotemporal behavior of DC-SIGN plays a crucial role not only on virus capture but also in clathrin-mediated endocytosis^29, 30^. We further proposed that Galectin-9 (Gal-9), a tandem repeat galectin with two heterologous CRDs^31^, and CD44, a heavily glycosylated protein and ligand of Gal-9 on T cells and NK cells^32, 33, 34^, might tune DC-SIGN-mediated clathrin-dependent endocytosis of viral particles^30^. However, a direct visualization of this process is still lacking due to the technical challenges of simultaneously combining SVT with multi-color SPT of DC-SIGN, CD44 and Gal-9.

Here we report on the implementation of a multi-color high-density single molecule imaging scheme based on quantum dot (QD) labeling to generate simultaneous, three-color high density maps (HiDenMaps) of DC-SIGN, CD44 and Gal-9 on the cell membrane of living primary imDCs isolated from healthy donors. Temporal autocorrelation analysis of HiDenMaps revealed a basal spatiotemporal inter-connection of these three proteins at the mesoscale, *prior* to virus engagement. We further extended the generation of HiDenMaps to four colors by including an additional fluorescence channel for GFP-tagged HIV-1 (eGFP-Gag) and SARS-CoV-2 virus like particles (VLPs). Remarkably, multi-color HiDenMaps allowed the systematic analysis of viral binding capture by the tripartite proteins in a multi-component fashion, clearly outcompeting with standard multi-color SPT in terms of throughput and efficiency. Our results show enhanced colocalization of HIV-1- and SARS-CoV-2-VLPs with the tripartite DC-SIGN, CD44 and Gal-9 nanoplatforms as compared to mock viruses. Importantly, we found that nanometer-scale colocalization of all three proteins enhanced DC-SIGN-mediated successful capture of HIV-1 and SARS-CoV-2-VLPs. Overall, multi-color HiDenMaps open the possibility for systematic studies of multi-molecular systems at the single molecule level in living cells with high throughput without compromising spatiotemporal resolution or single molecule detection sensitivity.

## Results

### Generation of multi-color HiDenMaps from high-density single molecule imaging

To perform multi-color single molecule imaging we designed a single molecule-sensitive inverted fluorescence microscope working in total internal reflection (TIR) configuration (Supplementary Text 1 and Fig. 1a). We immunolabeled DC-SIGN and CD44 with QD-single chain conjugates (QD655, QD705, respectively) and we tagged recombinant human Gal-9 with a biotin-streptavidin-QD (QD605) labeling scheme at a concentration of 30 nM (Methods and Supplementary Fig. 1). Virus-like particles (VLPs) were tagged with a green fluorescent protein (GFP-Gag). Importantly, considering the broad fluorescence excitation spectra of QDs and their overlap with that of GFP, we excited the four probes using a single diode-pumped solid-state 488 nm laser line. The four color fluorescence emission was collected through a Nikon CFI APO TIRF 60x objective and split into four optical paths so that each EM-CCD camera records two different spectral ranges (Fig. 1a). Importantly, the crosstalk between the different fluorescence channels ranged between 5-10% allowing multi-color imaging with minimal color crosstalk (see Supplementary Text 2 and Supplementary Fig. 2).

**Fig. 1:**
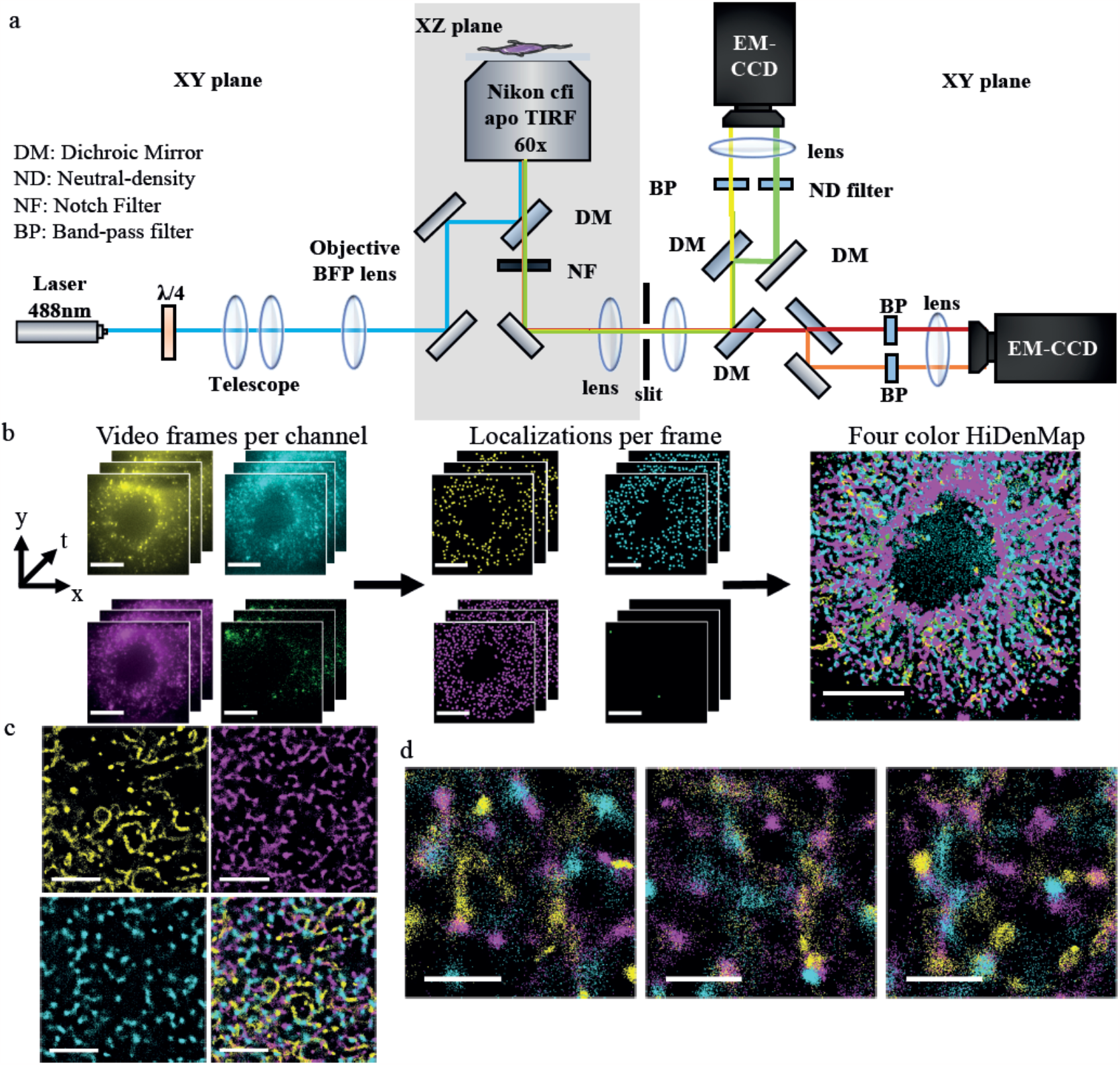
Generation of HiDenMaps from high-density multi-color single molecule imaging in living cells. **a**, Schematics of the custom-built four-color TIRF inverted microscope. The sample is illuminated by a 488 nm laser line focused on the back focal plane of the objective. The fluorescence emission is collected through the objective and split into four optical paths using dichroic mirrors and filtered using appropriate band-pass filters. Each emission is focused on two different regions of an EM-CCD camera, leading to a four-color detection scheme. **b**, Schematics on how four-color HiDenMaps are generated. From left to right: Video frames for each of the four channels. For each channel individual fluorescent molecules are localized at each frame of the video (middle panel). The localizations per frame and channel are then transformed using an affine transformation to correct for the aberration of the optical paths and collapsed into a single image (right panel). The localization precision of each individual QD resulted in 18.3 nm, 9.86 nm and 19.4 nm for QD605, QD655 and QD755, respectively, and 27.7 nm for the VLPs. Scale bars, 10 μm. (**c, d)**, Representative three-color HiDenMaps with an integration time of 30 seconds. **c**, DC-SIGN (yellow), CD44 (magenta), Gal-9 (cyan) and overlay of the three proteins (lower right panel). Scale bars, 10 μm. **d**, Representative enlarged overlay regions of the three proteins on different regions of the cell membrane. Scale bars, 1 μm.

To generate multi-color HiDenMaps from the recorded videos, we localized single molecules with the FIJI plugin Trackmate at each video frame of each channel and collapsed all the localizations into a single image using MATLAB^35^ (Fig. 1b). To account for variations in the expression level of each of the proteins due to cell-to-cell variability but also donor-to-donor variability, we normalized the number of localizations for every single HiDenMap (Supplementary Text 3). As a three-color example, we recorded a 90 seconds multi-color fluorescence video of DC-SIGN, CD44 and Gal-9 on the cell membrane of imDCs using a camera framerate of 30 Hz (Supplementary Videos 1-4). Remarkably, HiDenMaps generated by collapsing all the single molecule localizations for each of the channels during 30 seconds reveal that the three proteins dynamically explore the cell membrane in a rather heterogeneous fashion describing 2D patterns in their lateral diffusion (Supplementary Video 5 and Fig. 1c). Moreover, when visually observing in greater detail (3 μm x 3 μm zoom-in regions), it appears that the three proteins explore similar space in contiguous or even overlapping regions of the cell membrane (Supplementary Video 5 and Fig. 1d). These first observations hint to the existence of some type of correlated spatial (and/or temporal) organization of the tripartite proteins at the mesoscale (i.e., > 0.5 μm).

### HiDenMaps reveal that DC-SIGN, CD44 and Gal-9 explore the environment in an inter-connected manner

Since HiDenMaps encode both spatial and temporal information, we first extracted the relevant temporal scales by generating three-color HiDenMaps at different time windows, and acumulating the localizations during 2 seconds per window (see Methods). Similar to the previously generated maps built up with all the localizations (Fig. 1c,d), we observed contiguous spatial exploration of the tripartite proteins in all temporal windows (Fig. 2a), suggesting a coordinated diffusion over time. To quantify the degree of temporal correlation we used an overlapping sliding time-window of Δt = 500 ms to temporally accumulate localizations and computed the autocorrelation curves as function of time for the three proteins (Fig. 2b). The length of the window results from a trade-off of having to accumulate a sufficient number of localizations while retaining high-temporal resolution. The autocorrelation curves were best fitted using a double exponential function from which we extracted two characteristic decay time components (τ_1_, τ_2_) as well as the respective amplitudes, A_1_ and A_2_ (Methods and Supplementary Text 4). Notably, when we applied this type of analysis to the generated HiDenMaps of DC-SIGN, CD44 and Gal-9, we obtained comparable τ and A values for the three proteins (median values for τ_1_*≈* 3 s; τ_2_*≈* 30 s; A_1_*≈* 0.4 and A_2_*≈* 0.6) (Fig. 2c). The similarities in decay times and amplitudes of the autocorrelation curves indicate that the three proteins temporally explore their surrounding in a highly coordinated manner.

We next asked whether this “inter-connection” at the mesoscale is already established at the nanoscale. To tackle this question, we performed SPT of DC-SIGN, Gal-9 and CD44 using a reduced labeling density as compared to HiDenMaps. This approach allowed the reconnection of individual trajectories over nanoscale regions of the membrane at shorter temporal scales. By analyzing the time-averaged mean square displacement (Methods), we observed large differences in the distribution of the short-term diffusion values, *D*_1-4_, for DC-SIGN, CD44 and Gal-9 (Supplementary Fig. 3), indicating unrelated diffusion at the nanoscale. Overall, these results thus show that whereas at the nanoscale each protein diffuses and explores its nano-environment with different mobilities, a high degree of coordinated exploration exists over longer temporal and spatial scales.

**Fig. 2:**
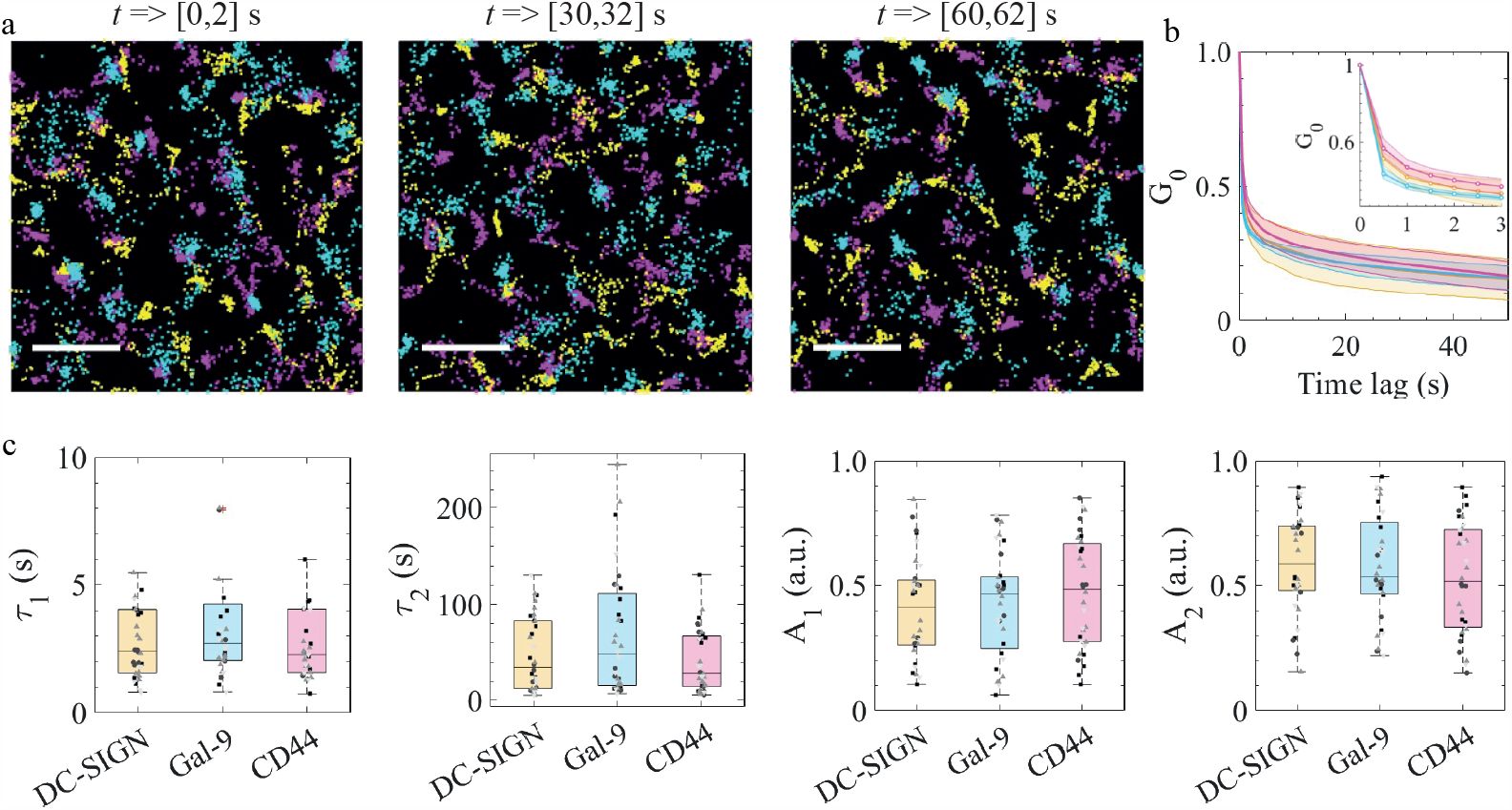
Temporal evolution of multi-color HiDenMaps for the tripartite proteins. **a**, Representative 2 seconds time windows of three-color HiDenMaps at different observation times. Yellow corresponds to DC-SIGN, magenta to CD44 and cyan to Gal-9. The imaging was performed at 30 Hz and the total acquisition was 90 s. Scale bars, 2 μm. **b**, Temporal autocorrelation decay curves for DC-SIGN, CD44 and Gal-9 with a Δt of 500 ms. The weighted line corresponds to the mean between donors and the shaded area the standard deviation between donors. The inset corresponds to the first three seconds of the autocorrelation curves. **c**, The parameters (τ_1_, τ_2_, A_1_, A_2_) fitted from the autocorrelation decays for DC-SIGN, Gal-9 and CD44. The whiskers represent the quartiles and the median value for each distribution. Each point in the scatter plot corresponds to a single cell. The data shown in **(b)** and **(c)** corresponds to three donors. For each donor we measured two samples, 10 cells per sample and roughly 8 ROIs per cell. Moreover, each donor was measured on a different day. The Kruskal-Wallis statistical test was performed and no statistical differences were found between the different proteins.

### Simultaneous three-color SPT and SVT reveals single HIV-1 VLP capture events by the tripartite proteins

To address if the spatiotemporal coordination of the tripartite proteins plays a role on the capture of HIV-1 VLPs, we first performed multi-color SPT of the three proteins in combination with SVT of a Gag-GFP-labeled HIV-1 VLP decorated with the BL4-3 envelope glycoprotein (Env), the ligand of DC-SIGN (see Methods). We focused on events where we detected the presence of HIV-1 VLPs together with the simultaneous occurrence of trajectories of the three proteins in the neighborhood of the HIV-1 VLPs, and followed their evolution in time (Fig. 3a). As expected from our previous results, tracking of the individual trajectories showed contiguous diffusion of the three proteins over time prior to HIV-1 VLP engagement (Fig. 3a,b, Supplementary Video 6). Notably, initial HIV-1 VLP capture in the close vicinity of the tripartite proteins occurred in an intermittent fashion, with multiple binding and unbinding events that finally resulted in a more stable binding and coincided with an increase in the spatial proximity of the three proteins (Fig. 3c). To better quantify these observations, we generated temporal windows of 2 seconds and overlaid the single molecule localizations of the three proteins and of the HIV-1 VLP into single maps (Fig. 3d, Supplementary Video 7). We then used a DBSCAN cluster algorithm^36^ to estimate the effective region explored by each of the proteins as function of time, prior and during HIV-1 VLP engagement (see Methods). This analysis showed a clear and gradual reduction in the cluster radius of the three different proteins during the capture event together with a concomitant and progressive increase in the number of HIV-1 VLP localizations being detected (Fig. 3e, Supplementary Video 7). Thus, these data reveal that HIV-1 VLP binding occurs on regions of the cell membrane already pre-enriched with the tripartite proteins and that their proximity is increased during viral engagement, enhancing in turn viral binding strength. To further strengthen these results we analyzed the number of localizations per protein cluster during viral capture. Indeed, while for DC-SIGN, CD44 and Gal-9 the number of localizations remained constant over time indicating stable engagement of the tripartite proteins during HIV-1 VLP capture, the number of HIV-1 VLP localizations increased over time, consistent with virus binding stabilization (Fig. 3f). As a whole, these results suggest that the spatiotemporal coordination of the tripartite proteins *prior* to virus engagement is an important step for the successful capture of HIV-1 VLPs, and significantly, that the binding strength of the virus is directly related to the spatiotemporal residence of the tripartite proteins at the virus-docking site.

**Fig. 3:**
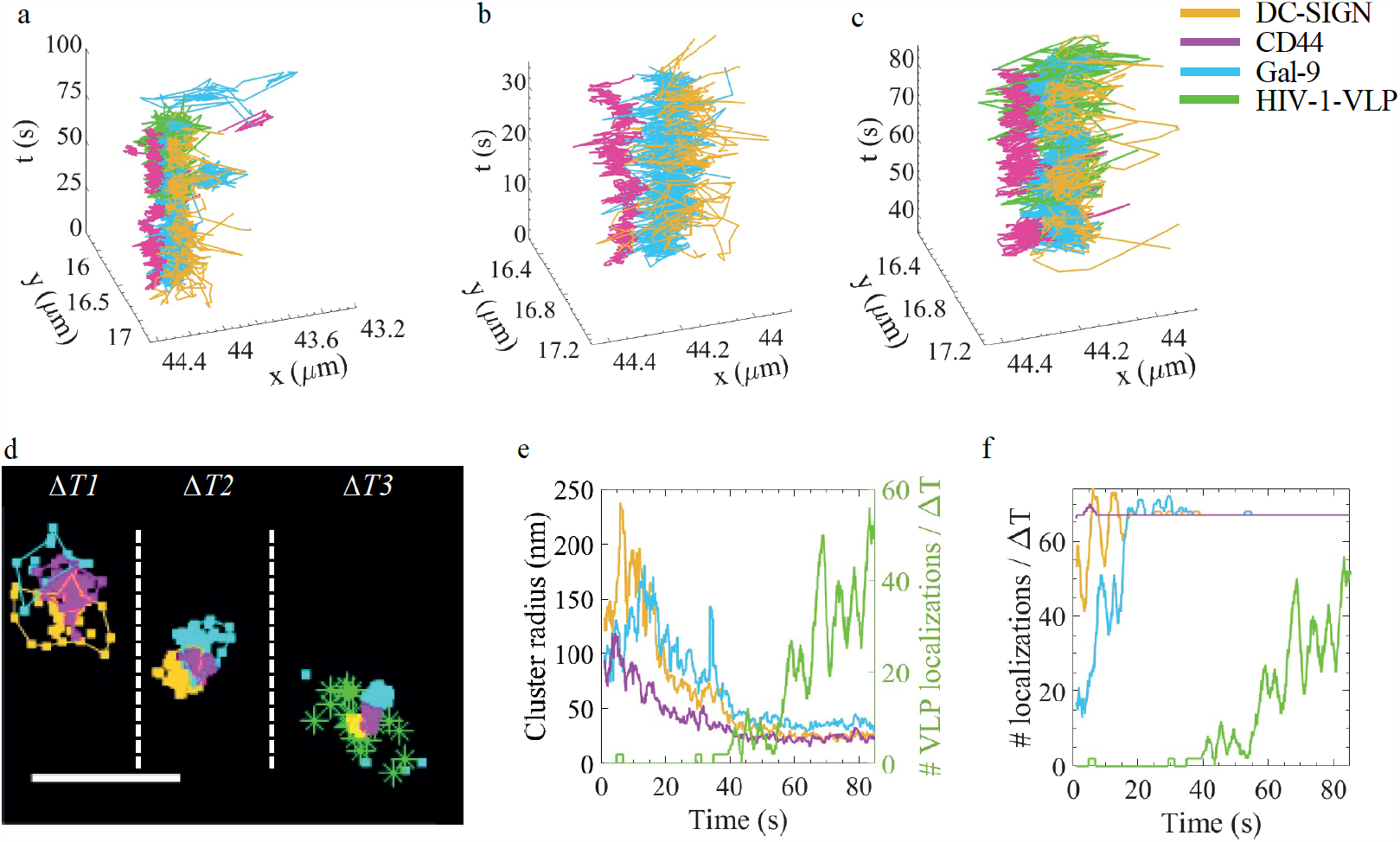
Temporal tracking of single HIV-1 VLP engagement by the tripartite proteins followed by simultaneous multi-color SPT and SVT. **a**, Single trajectories of DC-SIGN (yellow), CD44 (magenta), Gal-9 (cyan) and HIV-1 VLP (green) as function of time. **(b**,**c)**, Enlarged temporal windows of (**a**), prior to (**b**) and during VLP engagement (**c**). **d**, Snapshots of three representative temporal windows showing accumulated localizations (during 2 seconds) of the positions explored by DC-SIGN, CD44 and Gal-9, prior to (ΔT_1_, ΔT_2_) and during VLP engagement (ΔT_3_). The connecting lines between the symbols correspond to the identification and tracking of clusters of localizations for each of the proteins. The three proteins diffuse contiguous to each other prior to VLP engagement and their spatial proximity increases during VLP engagement. The three time-windows shown are: Δ*T*_1_ ∈ [2,4]*s*, Δ*T*_2_ ∈ [25,27]s and Δ*T*_X_ ∈ [70,72]*s*. Scale bar, 500 nm. **e**, Quantification of cluster radius (left axis) and the number of VLP localizations per time-window (2 seconds time-windows) (right axis) as function of time. **f**, Number of localizations per cluster and time-window for the three proteins and for the HIV-1 VLP (2 seconds time-windows). In (**e**,**f**), the data for the number of HIV-1 VLP localizations is the same.

### Four-color HiDenMaps reveal enhanced interactions of HIV-1 and SARS-CoV-2 VLPs with the tripartite protein nanoplatforms

To statistically enquire on the different spatiotemporal interactions between the tripartite proteins in the context of virus engagement we generated HiDenMaps for each of the three proteins and for the HIV-1 VLPs (Fig. 4a) and overlaid all the localizations in a single map (Fig. 4b). Visual inspection of such maps showed hotspots of enriched localizations for each of the proteins, forming distinct nanoplatforms on the cell membrane that spatially overlap with each other (white arrows in Fig. 4a,b). Remarkably, we observed a high degree of colocalization of multiple HIV-1 VLPs on regions enriched by the three protein nanoplatforms (white arrows in Fig. 4a,b). In strong contrast to standard SPT approaches where the probability of observing such events is vanishing small, HiDenMaps are able to uncover multiple virus-receptor interactions at the single molecule level, underscoring its effectiveness for monitoring multi-molecular interactions in real time.

**Fig. 4:**
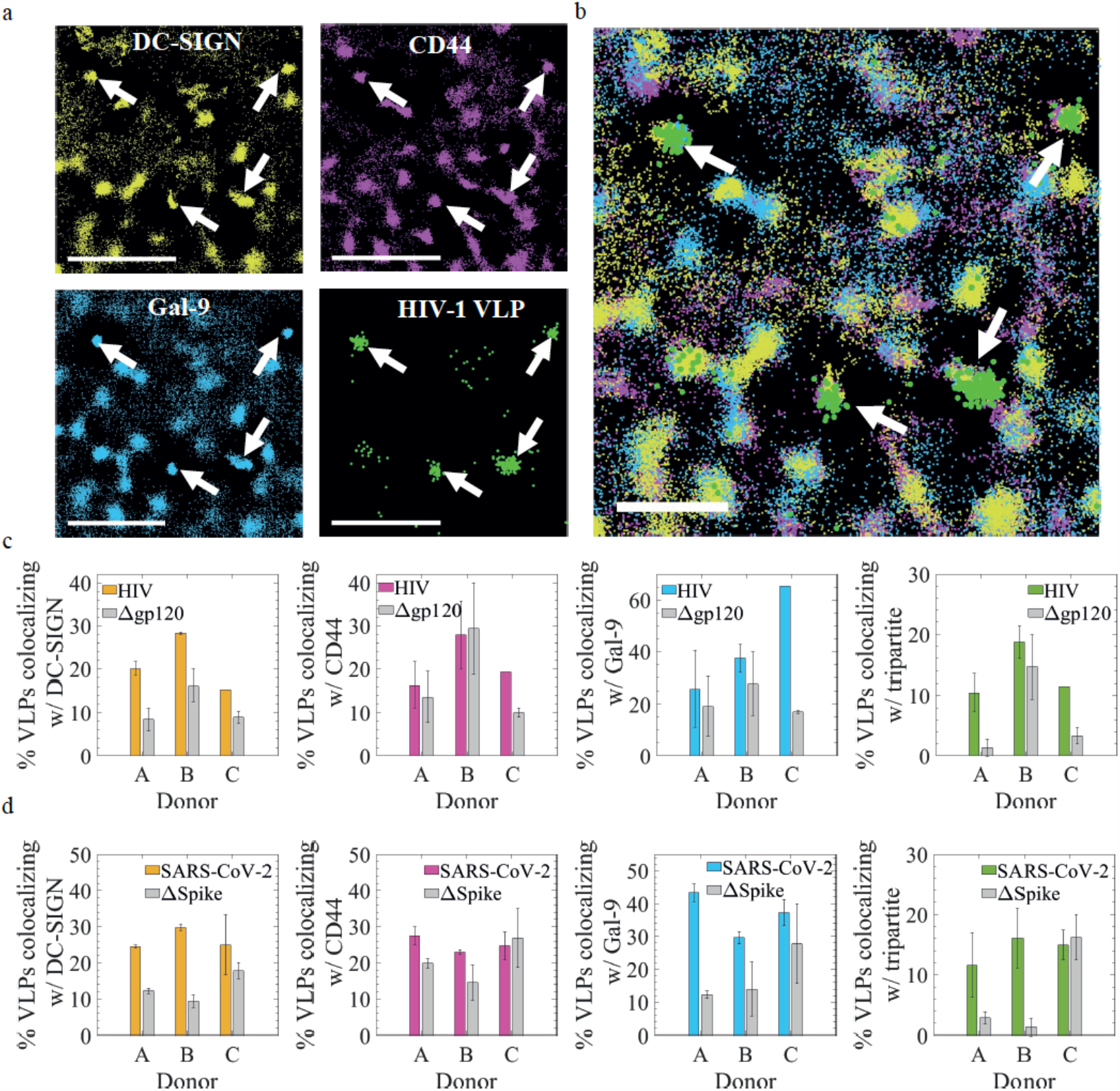
HIV-1 and SARS-CoV-2 VLPs exhibit enhanced spatiotemporal colocalization with the tripartite nanoplatforms. **a**, Representative four-color HiDenMaps of DC-SIGN (yellow), CD44 (magenta), Gal-9 (cyan) and HIV-1 VLP (green), generated from accumulating localizations during 50 seconds. White arrows in the different panels highlight multiple occurrence of simultaneous spatial colocalization of the tripartite proteins and the HIV-1 VLPs. Scale bars, 1 μm. **b**, Overlayed HiDenMap of the different panels shown in (**a**). **c**, Percentage of HIV-1 VLPs (colored bars) colocalizing with (from left to right): DC-SIGN, CD44, Gal-9 and with all three proteins, for three different donors investigated. Results obtained with the mock virus (Δgp120) are included as grey bars. **d**, Similar as (**c**) but for SARS-CoV-2 VLPs (colored bars) and mock virus (ΔSpike) (grey bars). Total number of HIV-1 VLPs detected: 64 for donor A, 53 for donor B and 26 for donor C. Total number of Δgp120 VLPs detected: 88 for donor A, 52 for donor B and 216 for donor C. Number of cells analyzed for the HIV-1 VLP experiments: 40 for donor A, 40 for donor B and 30 for donor C. Total number of SARS-CoV-2 VLPs detected: 132 for donor A, 74 for donor B and 87 for donor C. Total number of ΔSpike VLPs detected: 106 for donor A, 88 for donor B and 52 for donor C. Number of cells analyzed for the SARS-CoV-2 VLPs experiments: 40 for donor A, 40 for donor B and 40 for donor C. The bars represent the standard deviation between two independent experiments.

To quantify these observations we identified clusters of localizations for all the four different components and performed colocalization analysis between HIV-1 VLPs and the different proteins (Supplementary Text 5). We obtained a significant increased (15-30%) colocalization between DC-SIGN and HIV-1 VLPs as compared to mock VLPs in all donors analyzed (Fig. 4c, left panel) consistent with the fact that DC-SIGN is the main receptor for HIV-1 on imDCs^23, 30^. In the case of CD44 and Gal-9, only one donor (Donor C) exhibited increased colocalization with HIV-1 VLPs as compared to mock VLPs (Fig. 4c, middle panels). Notably, when analyzing the three proteins together, a remarkable high colocalization with HIV-1 VLPs above that of mock viruses was obtained in two out of the three donors analyzed (Fig. 4c, right panel). To further confirm that the measured colocalization between the tripartite proteins and HIV-1 VLPs is real and not the result of stochastic binding of the viruses to these regions, we also compared the experimental data to *in-silico* simulations of HIV-1 VLPs randomly landing on experimentally obtained HiDenMaps of the three different proteins on imDCs that have not been exposed to the VPLs (Supplementary Text 6). A low degree of colocalization between the tripartite proteins and the randomly distributed HIV-1 VLPs was obtained, comparable to that of the mock viruses (Supplementary Fig. 4). Together, these results reveal enhanced interaction of HIV-1-VLPs with pre-docking nanoplatforms formed by DC-SIGN, CD44 and Gal-9.

We then asked whether our methodology could be extended and generalized to the study of other virus-receptor interactions. Considering the urgence associated with the global COVID-19 pandemics, we applied our experimental settings to study SARS-CoV-2 VLPs capture by imDCs. Recent reports have identified DC-SIGN as a receptor for the SARS-CoV-2 virus, although the entry mechanisms remain to be elucidated^28, 37^. As DC-SIGN acts in concert with CD44 and Gal-9 to increase the binding of HIV-1 VLPs, we enquired whether a similar molecular mechanism could operate on the DC-SIGN-mediated binding of SARS-CoV-2. In order to generate SARS-CoV-2 VLPs, a plasmid encoding the S-protein ([D614G] the Wuhan variant with a point mutation) was expressed using the HIV-1 VLP backbone (Methods). The same mock virus without any receptor specific membrane protein, i.e., ΔSpike, was used as a negative control. Notably, more than 25% of SARS-CoV-2 VLPs colocalized with DC-SIGN in the three donors analyzed, significantly above that of the ΔSpike mutant (Fig. 4d, left panel), confirming that DC-SIGN is indeed a receptor for SARS-CoV-2 VLPs. Increased colocalization of SARS-CoV-2 VLPs with CD44 or Gal-9 was only clearly observed in one of the donors (Fig. 4d, middle panels). Remarkably and similar to HIV-1 VLPs, enhanced colocalization of SARS-CoV-2 VLPs with the tripartite proteins was observed in two out of the three donors, as compared to experiments performed using ΔSpike VLPs (Fig. 4d, right panel). Thus, these results strongly indicate that pre-docking nanoplatforms formed by the tripartite proteins favor the interaction of DC-SIGN, not only with HIV-1, but also importantly, with SARS-CoV-2.

### DC-SIGN, CD44 and Gal-9 pre-docking nanoplatforms enhance viral capture on the cell membrane of imDCs

Finally, we asked whether the enhanced interaction of the tripartite nanoplatforms with HIV-1 and SARS-CoV-2 VLPs results in increased viral capture. To address this question, the lateral behavior of individual viruses was analyzed and correlated with their interaction probability with the tripartite proteins. In other words, HiDenMaps of the different VLPs built up during 90 seconds observation times were generated and classified according to whether: (I) the VLP was engaged on the cell membrane before the acquisition started but vanished from the cell membrane before the end of the acquisition time. This behavior was classified as “vanishing” (Supplementary Video 8); (II) the VLP appeared and vanished during the acquisition time, being classified as “transient” (Supplementary Video 9); (III) the VLP appeared during the acquisition time and remained engaged on the cell membrane, being classified as “appearing” (Supplementary Video 10) and finally; (IV) the VLP remained engaged during the whole acquisition time. This behavior was classified as “persistent” (Supplementary Video 11). Representative examples of HiDenMaps from each category are shown in Supplementary Fig. 5. We then correlated these different VLPs behaviors to their *full* colocalization with the tripartite proteins, or to a *partial* colocalization, i.e., when the VLP colocalizes with only one or two of the tripartite proteins. Interestingly, in two out of the three donors we computed a higher percentage of appearing and persistent HIV-1 VLP events coinciding with a full colocalization of the three proteins (Fig. 5a, left), whereas a lower percentage of these events were obtained when the HIV-1 VLPs partially colocalized with only one or two of the other proteins (i.e., partial) (Fig. 5a, left). Since the residence time of the VLPs on the cell membrane likely reflects the strength of viral engagement, we further grouped transient and vanishing VLPs behaviors and classified them as unsuccessful viral capture events whereas appearing and persisting behaviors were classified as successful events. Remarkably, higher successful engagement of HIV-1 VLPs was observed when the three proteins fully colocalized with the virus in two of the three donors, as compared to VLPs that partially colocalized with only one or two of the proteins (Fig. 5a, right). These results imply that binding of HIV-1 VLPs by the three proteins increases the probability of virus capture on the cell membrane of imDCs. Moreover, successful viral capture by the tripartite proteins was higher for the HIV-1 VLPs as compared to VLPs lacking the envelope protein (BL4-3) for all three donors (Fig. 5b, HIV-1 *vs*. Δgp120), confirming once more the specificity of DC-SIGN in HIV-1 viral capture.

**Fig. 5:**
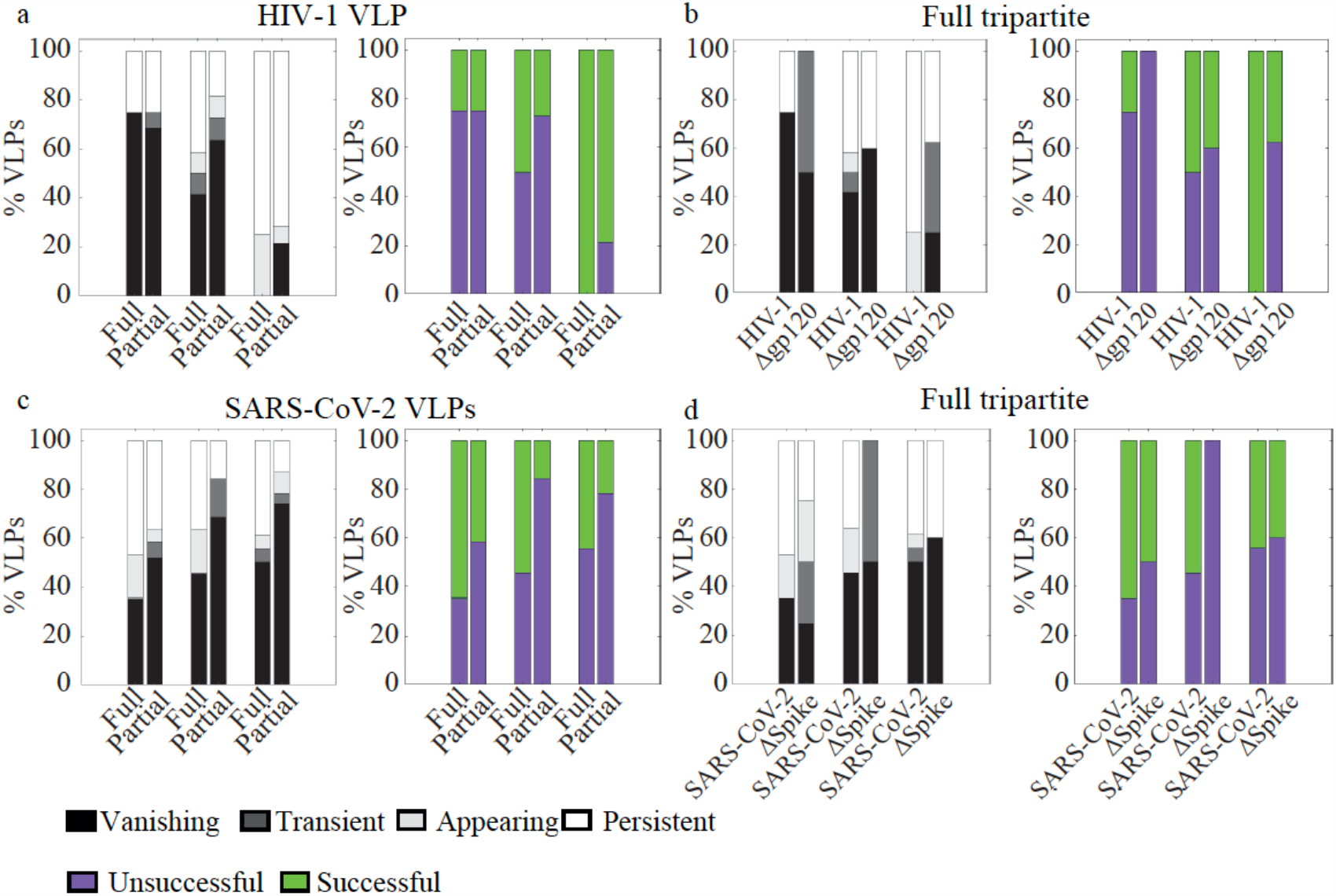
HIV-1 and SARS-CoV-2 VLPs show enhanced viral engagements on the cell membrane upon simultaneous binding to the tripartite nanoplatforms. **a**, left panel: Percentage of HIV-1 VLPs showing different engagement dynamics (vanishing, transient, appearing and persistent) when they fully colocalize with all three proteins (DC-SIGN, CD44 and Gal-9) i.e., denoted as “Full” in the plot, or with only a subset of the proteins, denoted as “Partial”. Right panel: Percentage of successful (green) and unsuccessful (violet) HIV-1 VLP engagements corresponding to the left panel. Data are shown for three different donors (from left to right: Donors A, B, C). **b**, left panel: Dynamic engagement for VLPs colocalizing with all three proteins, for HIV-1 VLP and the mock virus Δgp120 VLPs. Right panel: Percentage of successful (green) and unsuccessful (violet) VLP engagements corresponding to the left panel. **c**, Similar to (**a**) but for SARS-CoV-2 VLP. **d**, Similar to (**b**) but for SARS-CoV-2 VLP and the mock virus ΔSpike. Total number of HIV-1 VLPs detected: 64 for donor A, 53 for donor B and 26 for donor C. Total number of Δgp120 VLPs detected: 88 for donor A, 52 for donor B and 216 for donor C. Number of cells analyzed for the HIV-1 VLP experiments: 40 for donor A, 40 for donor B and 30 for donor C. Total number of SARS-CoV-2 VLPs detected: 132 for donor A, 74 for donor B and 87 for donor C. Total number of ΔSpike VLPs detected: 106 for donor A, 88 for donor B and 52 for donor C. Number of cells analyzed for the SARS-CoV-2 VLP experiments: 40 for donor A, 40 for donor B and 40 for donor C.

Finally, we performed the same type of analysis with SARS-CoV-2 VLPs and also found an increased successful engagement of SARS-CoV-2 VLPs with the three proteins together (Fig. 5c, full *vs*. partial) in all three donors. Moreover, the involvement of DC-SIGN in the processes of successful SARS-CoV-2 VLPs capture was also confirmed, as higher successful engagement events by the tripartite proteins were observed on the full VLPs as compared to mock viruses lacking the Spike protein (Fig. 5d, SARS-CoV-2 *vs*. ΔSpike). Altogether, these results robustly show that pre-formed docking nanoplatforms of DC-SIGN, CD44 and Gal-9 enhance viral capture on the cell membrane of imDCs.

## Discussion

The multi-color HiDenMap methodology described here represents a major step forward towards the understanding of multi-molecular interactions at the single molecule level in living cells. As an example, we have combined multi-color HiDenMaps of viral receptors on the cell membrane of imDCs with HiDenMaps of single viruses thereby visualizing multiple receptor-viral engagements in real time and at the single molecule level. Using this powerful methodology, we simultaneously captured the spatiotemporal organization of DC-SIGN, a viral receptor of HIV-1 and SARS-CoV-2 viruses in imDCs, and of CD44 together with Gal-9. Remarkably, we discovered that these three proteins dynamically explore the cell membrane in a highly coordinated and synchronous manner. Such a coordinated diffusion is likely orchestrated by the cortical actin cytoskeleton and by the transient interactions (either directly or indirectly) of any of these proteins to actin. Indeed, the temporal scales retrieved from our analysis (∼3 seconds and ∼30 seconds) are consistent with the cortical actin acting as a master regulator on the spatiotemporal organization of the cell membrane^38^. Moreover, CD44 has been proposed to act as a picket protein that links the underlying actin cytoskeleton and the extracellular milieu^39, 40^ and previous results from our group showed the dependence of Gal-9 on DC-SIGN diffusion and its lateral organization on the cell membrane^30^. With the new data provided by the simultaneous HiDenMaps of the tripartite proteins we now postulate that Gal-9 might serve as a linkage to connect DC-SIGN to CD44, so that the concerted motion of the tripartite proteins is ultimately guided by the direct interaction of CD44 to actin.

We further addressed the relevance of these tripartite nanoplatforms and their coordinated spatiotemporal diffusion on the capture of HIV-1 and SARS-CoV-2 VLPs by introducing an additional GFP-tagged VLP fluorescence channel, extending our setup to a four color configuration. In stark contrast to standard SPT experiments that suffer from extremely low throughput, HiDenMaps enabled the visualization of multiple virus-receptor interactions in a single experiment. We resolved, for the first time to our knowledge, individual interactions between VLPs and DC-SIGN and most importantly, uncovered the relevance of CD44 and Gal-9 forming basal nanoplatforms *prior* to virus engagement and fully overlapping with DC-SIGN. This coordinated spatiotemporal organization resulted crucial for the successful engagement of both HIV-1 and SARS-CoV-2 VLPs. Indeed, we found an increased successful engagement of HIV-1 VLPs when the viruses colocalized with the tripartite proteins indicating that pre-docking nanoplatforms of the three proteins promote a more efficient viral capture. We also applied our methodology to SARS-CoV-2 VLPs. Since it has been recently reported that DC-SIGN is a receptor for SARS-CoV-2 viruses^28, 37^, we hypothesized that SARS-CoV-2 capture could also be mediated by DC-SIGN/CD44/Gal-9 pre-docking nanoplatforms on imDCs. Indeed, our results showed increased binding of SARS-CoV-2 viruses to the tripartite nanoplatforms as compared to mock viruses. These results thus suggest a potential generalized mechanism of virus capture being mediated by DC-SIGN/CD44/Gal-9 nanoplatforms on the membrane of imDCs. The crucial involvement of CD44 and Gal-9 in this process could pave the way to novel anti-viral therapeutic strategies targeting any of these two proteins.

In summary, the new methodology reported here brings a major advance to the field by enabling the study of multi-molecular interactions at the single molecule level, in real time while retaining all the advantages of single molecule imaging methods. We have applied this methodology to follow the dynamics of individual viruses in combination with up to three different proteins on the cell membrane of primary cells but it can certainly be extended to the study of other multi-component processes in the cell. As such, our experimental approach opens a new door in the quantitative study of multi-molecular dynamic processes at relevant spatiotemporal scales with the potential of uncovering new phenomena that have remained inaccessible until now given the limitations of current single molecule imaging methods.

## Methods

### Primary cell culture

Human immature dendritic cells (imDCs) were obtained from peripheral blood mononuclear cells (PBMC) from HIV-1 seronegative donors using a Ficoll-Hypaque gradient (Alere Technologies AS). The monocyte population was selected by adherence on a T75cm^2^ flask for 1 hour. imDCs were obtained by culturing the monocytes in complete RPMI with 1.000 IU/ml GM-CSF (granulocyte-macrophage colony-stimulating factor) and IL-4 (interleukin-4) both from R&D for 6 days. The medium was replaced every two days with fresh GM-CSF and IL-4. Experiments were performed at day 6 from the monocyte extraction.

### Antibodies and reagents

Monoclonal mouse anti-human CD44 (Clone G44-26) and monoclonal mouse anti-CD209 (Clone DCN46) were obtained from BD Biosciences. Recombinant human Gal-9 protein (Cat. number 9064-GA) was obtained from R&D systems. SARS-CoV-2 Spike Protein (RBD) Chimeric Recombinant Rabbit Monoclonal Antibody (P05DHuRb) tagged with Alexa Fluor™ 647 was obtained from eBioscience. Streptavidin QDs (565, 605, 655 and 755) were obtained from Thermo Fisher scientific.

### Single chain antibody generation

All the experiments reported here have been performed using single chain antibodies to label DC-SIGN and CD44 in order to prevent antibody cross-linking (see Supplementary Fig. 1a for the general labeling strategy used for the generation of SPT and HiDenMaps). Both anti-human CD209 and CD44 single chain antibodies were generated using a similar protocol. First, the full chain antibodies were dialyzed using 10K dialysis devices (Thermo Scientific™ Slide-A-Lyzer™ MINI Dialysis Devices, 10K MWCO) against PBS for 8 hours at 4 ºC. Second, we concentrated the dialyzed full chain antibodies to a concentration of 1 mg/ml. We then reduced the antibodies using DTT (1,4-dithiotrheritol, Sigma Aldrich) at 1 mM, let the mixture to reduce at room temperature for 1 hour while rotating and after that dialyzed overnight against PBS using the 10K dialysis devices at 4 ºC. We stabilized the broken sulphide-bonds with Iodoacetamide at 20 mM. We let the mix at room temperature for 1 hour rotating gently and dialyzed to remove excess iodoacetamide overnight at 4 ºC. Supplementary Fig. 1b shows the electrophoresis gel for the generated single chain antibodies together with the full chain antibodies as a control.

### Biotinylation and conjugation to quantum dots

We performed the biotinylation of single chain antibodies and of Gal-9 with EZ-Link Sulfo-NHS-LC-Biotin (Thermo Fisher). For this, we added a 20x mol excess of biotin and let the mixture to shake for 1 hour in ice. Then, we dialyzed using 10K units overnight at 4 ºC to remove excess of non-reacting biotin. To conjugate the biotinylated single chains and Gal-9 to the QDs, we mixed equal ratios of single-chain/Gal-9 and QDs in 5x excess of free biotin. To obtain a target concentration of 300 nM of stock conjugates, we first mixed 300 nM of QDs with 1.5 μM biotin and then added 300 nM of single-chain/Gal-9. Importantly, to avoid artifacts due to crosslinking of the recombinant added Gal-9 proteins, titrations were performed until single molecules of the conjugate Gal-9/QD were detected.

### Labeling strategy

For SPT experiments, we used conjugates of Gal-9/QD565, α-CD44/QD655 and α-DC-SIGN/QD705 at a concentration of 1 nM. For the HiDenMap experiments, we increased their concentration to 30 nM.

### Pseudo-virus like particle generation

The plasmid pr8ΔEnv.2 was obtained from Addgene, Plasmid #12263). We generated VLPs as follows: 57 μL of Trans-IT reagent (Mirus) were added to 2 μg of pR8ΔEnv.2, 3 μg of Gag-eGFP, 1 μg of pcRev and 3 μg of BL4-3 Env to generate HIV-1 VLPs, or 0.5 μg of SARS-CoV-2-Spike [D614G] to generate SARS-CoV-2 VLPs. To generate mock viruses, no plasmid generating the Env or the Spike protein was added. We then added the mixture to 1.9 mL of OPTIMEM (Gibco) and incubated for 15 minutes at room temperature. We added the mixture to 18 mL of DMEM with FBS and L-Glut and without antibiotics. We added the medium to HEK-293T cells at 80-90% confluency and collected the supernatant at day 3. We first centrifuged briefly the supernatant (500 x g for 10 minutes) and filtered the supernatant through a 0.45 μm filter. We concentrated the supernatant with Lenti-X concentrator (Takara) following the manufacturer’s protocol and resuspend it in RPMI. Finally, the VLPs were aliquoted and kept under liquid nitrogen before storing them at – 80 ºC.

### Sample preparation for HiDenMap experiments

We plated ∼50.000 cells on either glass coverslips (#1) coated with PLL (20 ng/ml) for control cells or on 35 mm Glass bottom dish with 10 mm micro-well (#1, Cellvis) also coated with PLL. We seeded the cells for at least 1 hour in RPMI without FBS, L-Gluy or antibiotics. Labeling using the single chain-QDs was performed sequentially by diluting 1 μL of DC-SIGN/QD655 and 3 μL of CD44/QD705 in 46 μL of PBS with 6% BSA. We incubated the conjugates and the cells for 5 min. After washing 3 times in RPMI, we took 5 μL of Gal-9/QD605 and 45 μL of PBS diluted in 6% BSA and incubated for 5 min. Importantly and to avoid removal of the Gal-9 conjugate, we washed only once with RPMI. We then added RPMI to perform the imaging. For the experiments with VLPs (HIV-1, SARS-CoV-2 or mocks) we added the VLPs defrosted and added 10 ng/ml of LPS. The mixture was added to the cells and immediately imaged at 37 ºC on the basal membrane.

### Spatiotemporal autocorrelation decay curves

For the analysis on the spatiotemporal autocorrelation functions, we took the HiDenMap generated over 90 seconds of acquisition and defined regions of interest (ROIs) of 5-by-5 μm. Then, we took all the localizations for the different HiDenMaps of the different proteins and applied an overlapping sliding time window, Δt = 500 ms to temporally separate the localizations. We then compute each autocorrelation curve as follows:

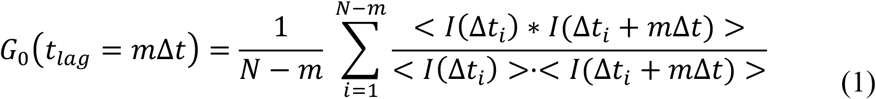

Here, I(Δt_i_) refers to the image of the i-th temporal window. We then normalized the curve to the first point (t_lag_ = 0 s). Finally, we fitted the decay curves from the second point onwards until t_lag_ = 50 s. We found that the best goodness of the fit consisted on a double exponential decay with a constant term:

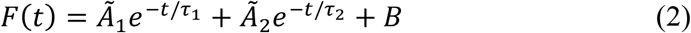

Finally, once the fitting was performed, we rescaled the amplitudes of the exponential decays as follows to ascertain for the relative weight of the two fitted exponentials:

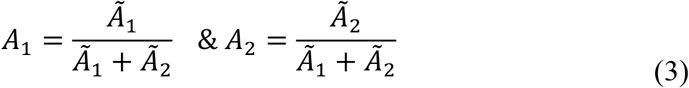

See Supplementary Text 3 for more detailed explanations over the temporal window chosen and the autocorrelation fitting procedures.

### SPT experiments and diffusion analysis from individual trajectories of DC-SIGN, CD44 and Gal-9

We performed three-color SPT by labeling CD44 and DC-SIGN with conjugated single-chain antibodies with QD655 and QD705 (respectively) and we conjugated recombinant Gal-9 biotinylated with QD565. We performed the imaging at 30Hz and acquired videos of 90s. We then performed the tracking using Trackmate (FIJI) and filtered those trajectories shorter than 50 frames (1.5 s). We computed the short-term diffusion coefficient, D_1-4_ by fitting the mean-square displacement (MSD) slope for the first 4 time points, in a similar way as described in^41^.

### Determination of clusters of localization and their radius using DBSCAN

To quantify the cluster areas of Gal-9, CD44 and DC-SIGN, 1x1 um ROIs of the VLPs were selected as the reference ROI. Then a DBSCAN analysis with ε=300 nm and the minimum number of points of 10 for Gal-9 and 20 for DC-SIGN and CD44 was computed. To calculate the cluster radius, the MATLAB function polyarea was used.

### Software

For the SPT detection and linking of trajectories we have used ImageJ’s FIJI plugin Trackmate^35^. All the data have been analyzed using MATLAB R2020a.

## Acknowledgments

We would like to thank Carlo Manzo and Felix Campelo for fruitful discussions. The research leading to these results has received funding from the European Commission H2020 Program under grant agreement ERC Adv788546 (NANO-MEMEC) (to M.F.G.-P.), ERC-2019-CoG-863869 (FUSION) (to S.P.-P.), The Chan-Zuckerberg Initiative “Multicolor single molecule tracking with lifetime imaging” (2023-321188 to S.P.-P.), Government of Spain (Severo Ochoa CEX2019-000910-S, State Research Agency (AEI) (PID2020-113068RB-I00 / 10.13039/501100011033 (to M.F.G.-P.) and JdC-IJCI-2017-33160 (to J.A.T.-P.), Fundació CELLEX (Barcelona), Fundació Mir-Puig and the Generalitat de Catalunya through the CERCA program and AGAUR (Grant No. 2021 SGR01450 M.F.G.-P.). N.M. acknowledges funding from the European Union H2020 under the Marie Sklodowska-Curie grant 754558-PREBIST.

## Author Contributions

J.A.T.-P. and M.F.G.-P. designed research; N.M., J.A.T.-P., E.G.-M., J.A.-C. performed experiments; N.M. designed software tools and performed analysis of the data; I.C.-A., S.P.-P contributed with new reagents and designed the composition of the VLPs; S.P.-P., J.A.T.-P., M.F.G.-P., supervised research; N.M., J.A.T.-P., M.F.G.-P. wrote the manuscript; all authors read and provided feedback to the manuscript.

The authors declare no competing interest

## References

1. Haynes, B.F. et al. Strategies for HIV-1 vaccines that induce broadly neutralizing antibodies. Nat Rev Immunol 23, 142–158 (2023).

2. Mahajan, S., Choudhary, S., Kumar, P. & Tomar, S. Antiviral strategies targeting host factors and mechanisms obliging +ssRNA viral pathogens. Bioorg Med Chem 46, 116356 (2021).

3. Pollard, A.J. & Bijker, E.M. A guide to vaccinology: from basic principles to new developments. Nat Rev Immunol 21, 83–100 (2021).

4. Rahman, M.M., Masum, M.H.U., Wajed, S. & Talukder, A. A comprehensive review on COVID-19 vaccines: development, effectiveness, adverse effects, distribution and challenges. Virusdisease 33, 1–22 (2022).

5. Johnson, C., Exell, J., Lin, Y., Aguilar, J. & Welsher, K.D. Capturing the start point of the virus-cell interaction with high-speed 3D single-virus tracking. Nat Methods 19, 1642–1652 (2022).

6. Lakadamyali, M., Rust, M.J., Babcock, H.P. & Zhuang, X. Visualizing infection of individual influenza viruses. Proc Natl Acad Sci U S A 100, 9280–9285 (2003).

7. Liu, S.L. et al. Single-Virus Tracking: From Imaging Methodologies to Virological Applications. Chem Rev 120, 1936–1979 (2020).

8. Seisenberger, G. et al. Real-time single-molecule imaging of the infection pathway of an adeno-associated virus. Science 294, 1929–1932 (2001).

9. Coomer, C.A. et al. Single-cell glycolytic activity regulates membrane tension and HIV-1 fusion. PLoS Pathog 16, e1008359 (2020).

10. Coller, K.E. et al. RNA interference and single particle tracking analysis of hepatitis C virus endocytosis. PLoS Pathog 5, e1000702 (2009).

11. Cureton, D.K., Harbison, C.E., Cocucci, E., Parrish, C.R. & Kirchhausen, T. Limited transferrin receptor clustering allows rapid diffusion of canine parvovirus into clathrin endocytic structures. J Virol 86, 5330–5340 (2012).

12. Ewers, H. et al. Single-particle tracking of murine polyoma virus-like particles on live cells and artificial membranes. Proc Natl Acad Sci U S A 102, 15110–15115 (2005).

13. Pelkmans, L., Kartenbeck, J. & Helenius, A. Caveolar endocytosis of simian virus 40 reveals a new two-step vesicular-transport pathway to the ER. Nat Cell Biol 3, 473–483 (2001).

14. Rust, M.J., Lakadamyali, M., Zhang, F. & Zhuang, X. Assembly of endocytic machinery around individual influenza viruses during viral entry. Nat Struct Mol Biol 11, 567–573 (2004).

15. van der Schaar, H.M. et al. Dissecting the cell entry pathway of dengue virus by single-particle tracking in living cells. PLoS Pathog 4, e1000244 (2008).

16. Schelhaas, M. et al. Human papillomavirus type 16 entry: retrograde cell surface transport along actin-rich protrusions. PLoS Pathog 4, e1000148 (2008).

17. Sun, E.Z. et al. Real-Time Dissection of Distinct Dynamin-Dependent Endocytic Routes of Influenza A Virus by Quantum Dot-Based Single-Virus Tracking. ACS Nano 11, 4395–4406 (2017).

18. Xu, H. et al. Real-time Imaging of Rabies Virus Entry into Living Vero cells. Sci Rep 5, 11753 (2015).

19. Zhang, L.J. et al. A “Driver Switchover” Mechanism of Influenza Virus Transport from Microfilaments to Microtubules. ACS Nano 12, 474–484 (2018).

20. Banchereau, J. & Steinman, R.M. Dendritic cells and the control of immunity. Nature 392, 245–252 (1998).

21. Cabeza-Cabrerizo, M., Cardoso, A., Minutti, C.M., Pereira da Costa, M. & Reis e Sousa, C. Dendritic Cells Revisited. Annu Rev Immunol 39, 131–166 (2021).

22. Zanna, M.Y. et al. Review of Dendritic Cells, Their Role in Clinical Immunology, and Distribution in Various Animal Species. Int J Mol Sci 22 (2021).

23. Geijtenbeek, T.B. et al. DC-SIGN, a dendritic cell-specific HIV-1-binding protein that enhances trans-infection of T cells. Cell 100, 587–597 (2000).

24. Harrison, S.C. Viral membrane fusion. Virology 479-480, 498–507 (2015).

25. Kwon, D.S., Gregorio, G., Bitton, N., Hendrickson, W.A. & Littman, D.R. DCSIGN-mediated internalization of HIV is required for trans-enhancement of T cell infection. Immunity 16, 135–144 (2002).

26. Cambi, A., Beeren, I., Joosten, B., Fransen, J.A. & Figdor, C.G. The C-type lectin DC-SIGN internalizes soluble antigens and HIV-1 virions via a clathrin-dependent mechanism. Eur J Immunol 39, 1923–1928 (2009).

27. Tacken, P.J. et al. Targeting DC-SIGN via its neck region leads to prolonged antigen residence in early endosomes, delayed lysosomal degradation, and crosspresentation. Blood 118, 4111–4119 (2011).

28. Amraei, R. et al. CD209L/L-SIGN and CD209/DC-SIGN Act as Receptors for SARS-CoV-2. ACS Cent Sci 7, 1156–1165 (2021).

29. Manzo, C. et al. The neck region of the C-type lectin DC-SIGN regulates its surface spatiotemporal organization and virus-binding capacity on antigenpresenting cells. J Biol Chem 287, 38946–38955 (2012).

30. Torreno-Pina, J.A. et al. Enhanced receptor-clathrin interactions induced by Nglycan-mediated membrane micropatterning. Proc Natl Acad Sci U S A 111, 11037–11042 (2014).

31. Moar, P. & Tandon, R. Galectin-9 as a biomarker of disease severity. Cellular immunology 361, 104287 (2021).

32. Ponta, H., Sherman, L. & Herrlich, P.A. CD44: from adhesion molecules to signalling regulators. Nat Rev Mol Cell Biol 4, 33–45 (2003).

33. Rahmati, A., Bigam, S. & Elahi, S. Galectin-9 promotes natural killer cells activity via interaction with CD44. Front Immunol 14, 1131379 (2023).

34. Wu, C. et al. Galectin-9-CD44 interaction enhances stability and function of adaptive regulatory T cells. Immunity 41, 270–282 (2014).

35. Tinevez, J.Y. et al. TrackMate: An open and extensible platform for singleparticle tracking. Methods 115, 80–90 (2017).

36. Ester, M., Kriegel, H.-P., Sander, J. & Xu, X. A density-based algorithm for discovering clusters in large spatial databases with noise. kdd; 1996; 1996. p. 226–231.

37. Lim, S., Zhang, M. & Chang, T.L. ACE2-Independent Alternative Receptors for SARS-CoV-2. Viruses 14 (2022).

38. Mateos, N. et al. High-density single-molecule maps reveal transient membrane receptor interactions within a dynamically varying environment. arXiv preprint arXiv:2307.07334 (2023).

39. Freeman, S.A. et al. Transmembrane Pickets Connect Cyto- and Pericellular Skeletons Forming Barriers to Receptor Engagement. Cell 172, 305–317 e310 (2018).

40. Sil, P. et al. Dynamic actin-mediated nano-scale clustering of CD44 regulates its meso-scale organization at the plasma membrane. Molecular Biology of the Cell 31, 561–579 (2020).

41. Munoz-Gil, G. et al. Stochastic particle unbinding modulates growth dynamics and size of transcription factor condensates in living cells. Proc Natl Acad Sci U S A 119, e2200667119 (2022).

