## Supplementary Information for "Early steps of multi-receptor viral interactions dissected by high-density, multi-color single molecule mapping in living cells"

[*] Corresponding authors:

**Supplementary Text 1: Four-color single molecule sensitive microscope.** The optical setup is built around an inverted Nikon Ti-U microscope with a total internal reflection (TIR) illumination configuration (Fig. 1a). The excitation light consists on a single diode-pumped solid-state 488 nm laser line which is made circularly polarized using a quarter waveplate. The beam diameter is then increased using a telescope and ultimately focused at the back focal plane of a Nikon CFI APO TIRF 60x objective. In order to obtain the TIR excitation, the beam is accurately shifted towards the edge of the objective using a mirror at the entrance of the inverted microscope. The emitted fluorescence light from the sample is collected through the same objective and the excitation light is filtered out using a notch-filter (NF) at 488nm. After creating the image using a slit, the emission light is split into four optical paths using appropriated dichroic mirrors and filtered with band-pass (BP) filters. Each emission is then focused on two different regions of an EM-CCD camera. The use of two EM-CCD cameras leads to a four-color detection scheme. To filter the emission from the QDs we use BP filters centered at 607 nm, 640 nm and 725 nm with a bandwidth of 30 nm. In the light path of GFP, we used a neutral density (ND) filter to attenuate any cross-talk signal from QD605. When single QDs or GFP-VLPs are placed on the set-up and measured separately, the crosstalk between channels ranges from 5 to 10% in intensity (see Supplementary Fig. 2 and Supplementary Text 2). Moreover, the crosstalk is even lower when the three different QDs and the GFP-VLPs are imaged simultaneously. This is due to the fact that the threshold of each intensity channel is increased when imaging the four channels.

The field of view (FOV) of each channel is 512x256 pixel with a pixel size of 119 nm making a total of 60-by-30 µm imaging areas. The frame rate and the total acquisition time used for all the experiments is 30 Hz and 90 seconds, respectively. The localization precision, estimated as the spread of single localizations from the mean value, of the different QDs used was 18.3 nm, 9.86 nm and 19.4 nm for QD605, QD655 and QD755, respectively, and 27.7 nm for the VLPs. In order to correct for optical aberrations of the four different fluorescence emission channels, we used multi-color TetraSpeck microspheres (0.1 µm) whose fluorescence emission was recorded simultaneously in all the channels. We then applied an affine transformation of the four channels having the GFP channel as reference. As such, we obtained an affine transformation error of 85 nm, 96 nm and 80 nm for the channels of DC-SIGN, Gal-9 and CD44, respectively.

**Supplementary Text 2: Crosstalk correction.** To assess the degree of crosstalk in the different fluorescence channels we imaged separately each individual channel with its corresponding QD or GFP-VLP. After localizing individual QDs or GFP-VLPs, their fluorescence signal was subtracted from the fluorescence background of the channel. The fluorescence background of each channel was extracted from a 5x5 pixel area without any detected QD or GFP-VLP. Then, the background corrected fluorescence signal was compared and quantified with the other three different channels yielding only a 5-10% crosstalk (Supplementary Fig. 2).

**Supplementary Text 3: Normalization of the localizations of HiDenMaps.** Since the expression level of each protein vary not only from cell-to-cell but also from donor-to-donor, we implemented an automated approach to normalize the number of localizations per region of interest (ROI). First, we defined ROIs of 12x12 pixel areas from each individual HiDenMap. Then, we generated a distribution of the total localizations of each protein within the 12x12 pixel area and calculated the mean value averaged through all the ROIs. We multiply this mean value by 0.65 and the resulting value was the reference number of localizations to take into account for each channel. We employ the value 0.65 as it was an optimal compromise between having a sufficient amount of ROIs containing the minimum number of localizations for our analysis. Indeed, if a ROI had a lower number of localizations than the reference value, the ROI was excluded from further analysis. On the other hand, if the ROI had a higher number of localizations than the reference value, localizations were stochastically excluded until reaching the reference value.

**Supplementary Text 4: Spatiotemporal autocorrelation decay curves.** As mentioned in the main text, we applied an overlapping sliding time-window, $\Delta t$, of 500 ms to temporally cumulate the localizations. Given the typical number of localizations contained in each ROI of the HiDenMap to be autocorrelated, we estimated that 500 ms was the best compromise between having a sufficient number of localizations (at least ~100 localizations per ROI in $\Delta t$) and generating a reliable autocorrelation decay curve. Since we normalize the number of localizations per each ROI for each protein (DC-SIGN, CD44, Gal-9), using shorter time windows would prevent the generation of reliable decay curves due to a lower number of localizations. Thus, a time window of 500 ms resulted optimal, taking into account the integration time and a consistent decay curve at each random sampling of localizations.

The autocorrelation curves for each given ROI were best fitted using a double exponential decay with a constant term:

$$F(t) = \tilde{A}_1 e^{-t/\tau_1} + \tilde{A}_2 e^{-t/\tau_2} + B \quad (1)$$

We performed the fitting in MATLAB's Curve Fitting Tool setting the bounds of $\widetilde{A_1}$, $\widetilde{A_2}$ and $B \in [0,1]$ and $\tau_1$ and $\tau_2 \in [0, \infty)$. We chose '0.5' as the starting point for all the fitted variables. The rest of parameters used for the fittings are summarized in the following table:

| **Algorithm** | **Trust-Region** |
|---|---|
| Robust | On |
| Maximum iterations | $10^5$ |
| Maximum number of evaluations | $10^5$ |
| Minimum change in coefficients | $10^{-8}$ |
| Maximum change in coefficients | $10^{-2}$ |
| Termination tolerance on model value | $10^{-16}$ |
| Termination tolerance on coefficient values | $10^{-16}$ |

**Supplementary Text 5**: **Multi-color colocalization algorithm.** To quantify the degree of colocalization between the VLPs and the proteins of interest (DC-SIGN, CD44 and Gal-9), we took the HiDenMaps of the VLP as the reference channel. We first defined VLP clusters by running a DBSCAN algorithm with parameters MinPts = 10 and ε = 150 nm in the VLP HiDenMap[1]. Once the VLP clusters are detected, we defined these clusters as the region of interest (ROI). If the other three proteins visit this ROI for the same amount of time as the VLP, we consider it as a full colocalization event (Fig. 4c,d and denoted as "Full" in Fig. 5). If only one or two proteins visit this ROI, we consider it as a partial colocalization event (Fig. 4c,d and denoted as "Partial" in Fig. 5). Importantly, we also consider a temporal correlation between these colocalization events, i.e., the proteins must colocalize for at least 10 consecutive frames and 95% of the localizations of each protein have to be detected inside the ROI within the temporal window.

**Supplementary Text 6: *In silico*-simulations of stochastic viral landings on the cell membrane.** To perform the *in-silico* simulations, we generated experimental HiDenMaps for each of the proteins of interest (DC-SIGN, CD44 and Gal-9) on imDCs that have been exposed to the VLPs or in control imDCs (i.e., non-exposed to VLPs). In the case of cells that have been exposed to the VLPs, we also generated the corresponding VLP-HiDenMap and extracted clusters of localizations of the VLPs. We then positioned the clusters of localizations of the VLPs randomly on the HiDenMaps of the control cells and calculated the degree of colocalization between the VLP clusters of localizations and each of the three proteins or to the tripartite proteins (Supplementary Fig. 4). Importantly, we performed this analysis on multiple regions of the cell membrane to minimize potential artifacts in the colocalization degree due to different expression levels or labeling efficiencies of the proteins.

**Supplementary Figures & legends:**

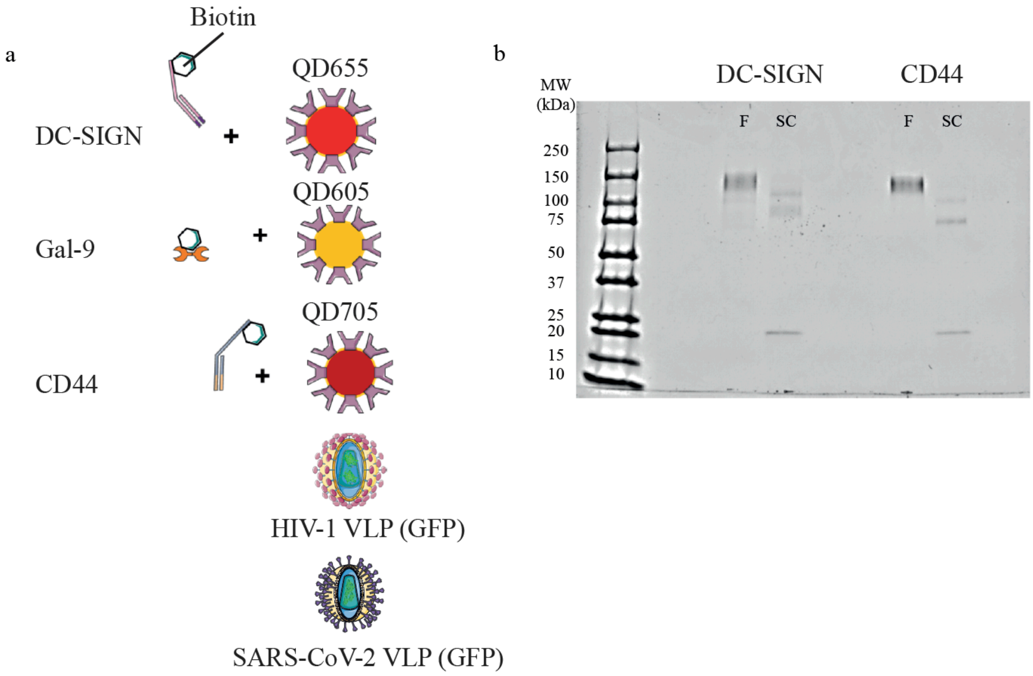


### SUPPLEMENTARY FIGURE 1

**Supplementary Fig. 1: Labeling strategy for HiDenMap experiments. a,** DC-SIGN and CD44 are labeled using single chain antibodies biotinylated and conjugated with streptavidin-coated QDs. We use biotinylated recombinant human Gal-9 conjugated with streptavidin-coated QD. VLPs were tagged using GFP (GFP-Gag). **b,** Electrophoresis gel showing the bands for full (F) and single chains (SC).

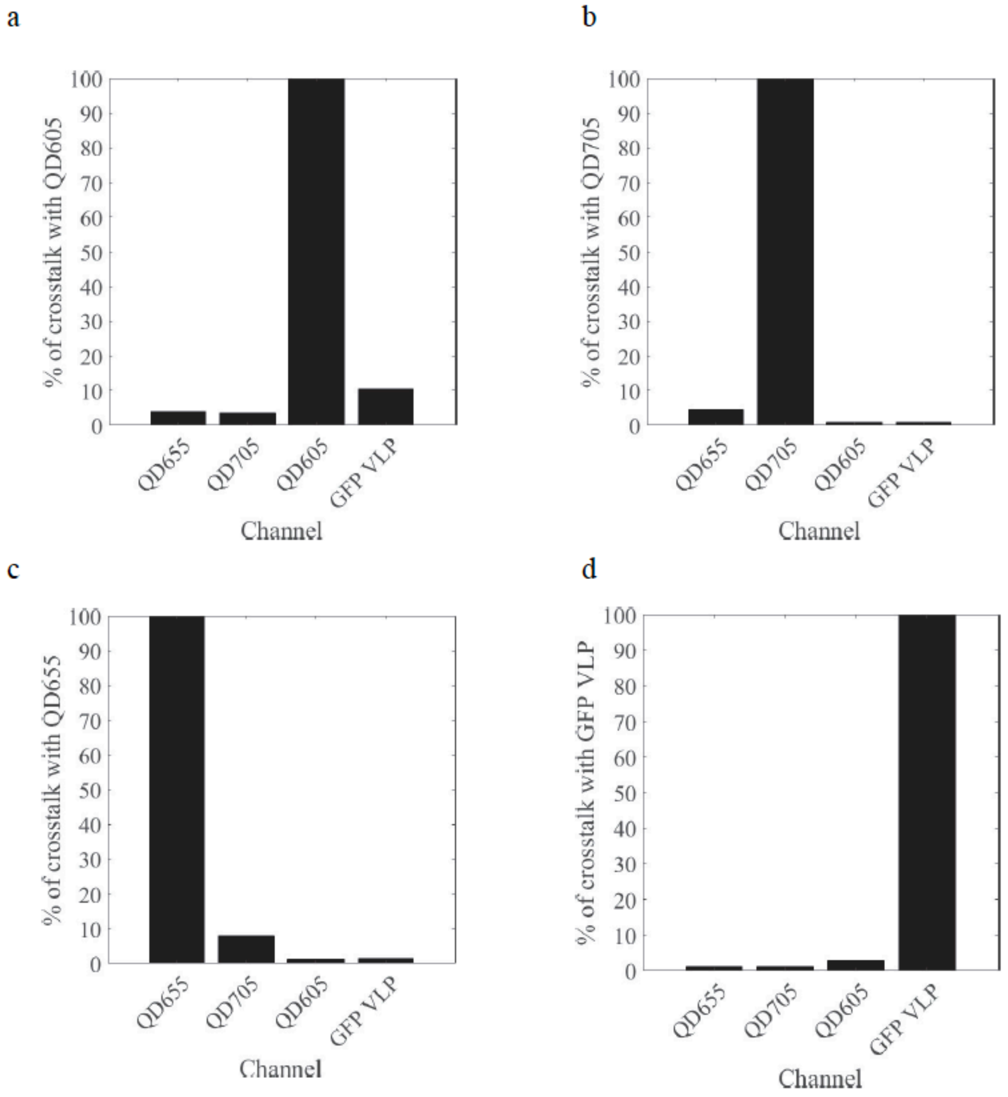


### SUPPLEMENTARY FIGURE 2

**Supplementary Fig. 2: Evaluation of the spectral crosstalk between the different channels.** Crosstalk of the four different fluorescence channels: QD605 (a), QD705 (b), QD655 (c) and GFP-VLP (d), with respect to the three remaining channels. The maximum crosstalk is ca. 10%.

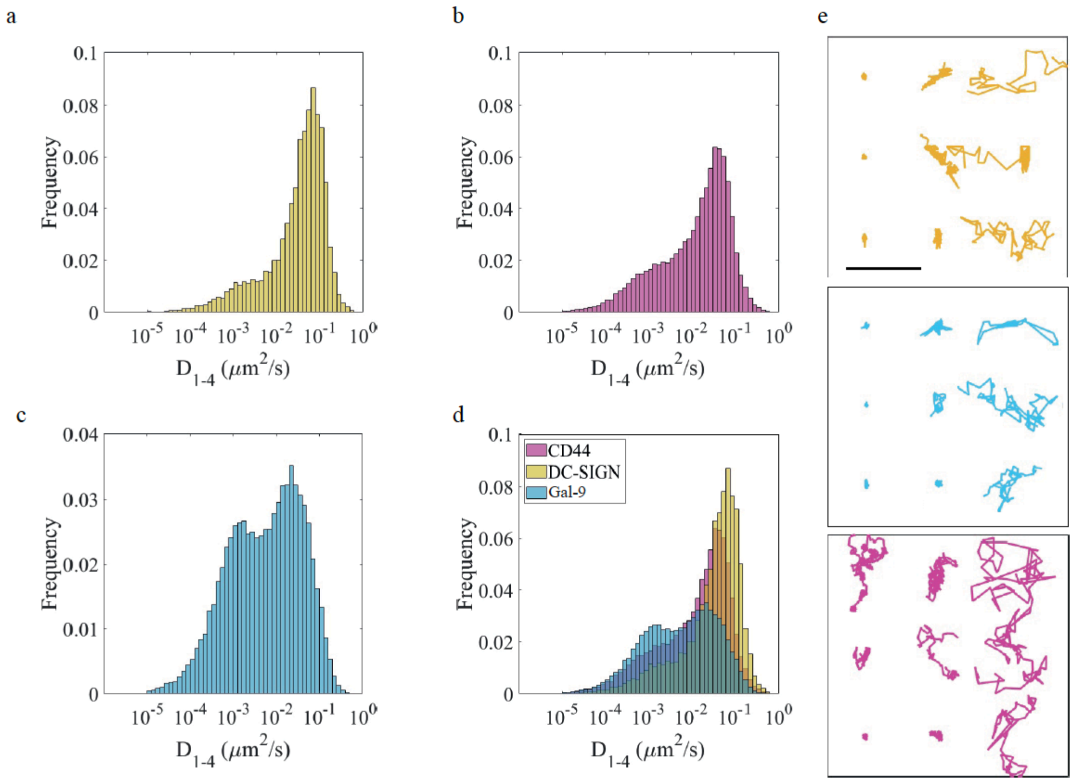


#### SUPPLEMENTARY FIGURE 3

**Supplementary Fig. 3: SPT experiments of the tripartite proteins show different diffusion behavior at the nanoscale. (a-d)** Histograms of the distribution of $D_{1\text{-}4}$ values for **a,** DC-SIGN, **b,** CD44 **c,** Gal-9 and **d,** the combination of the three different histograms. **e,** representative single molecule trajectories of the three proteins: DC-SIGN (yellow), Gal-9 (cyan) and CD44 (magenta). Scale bar, 500 nm.

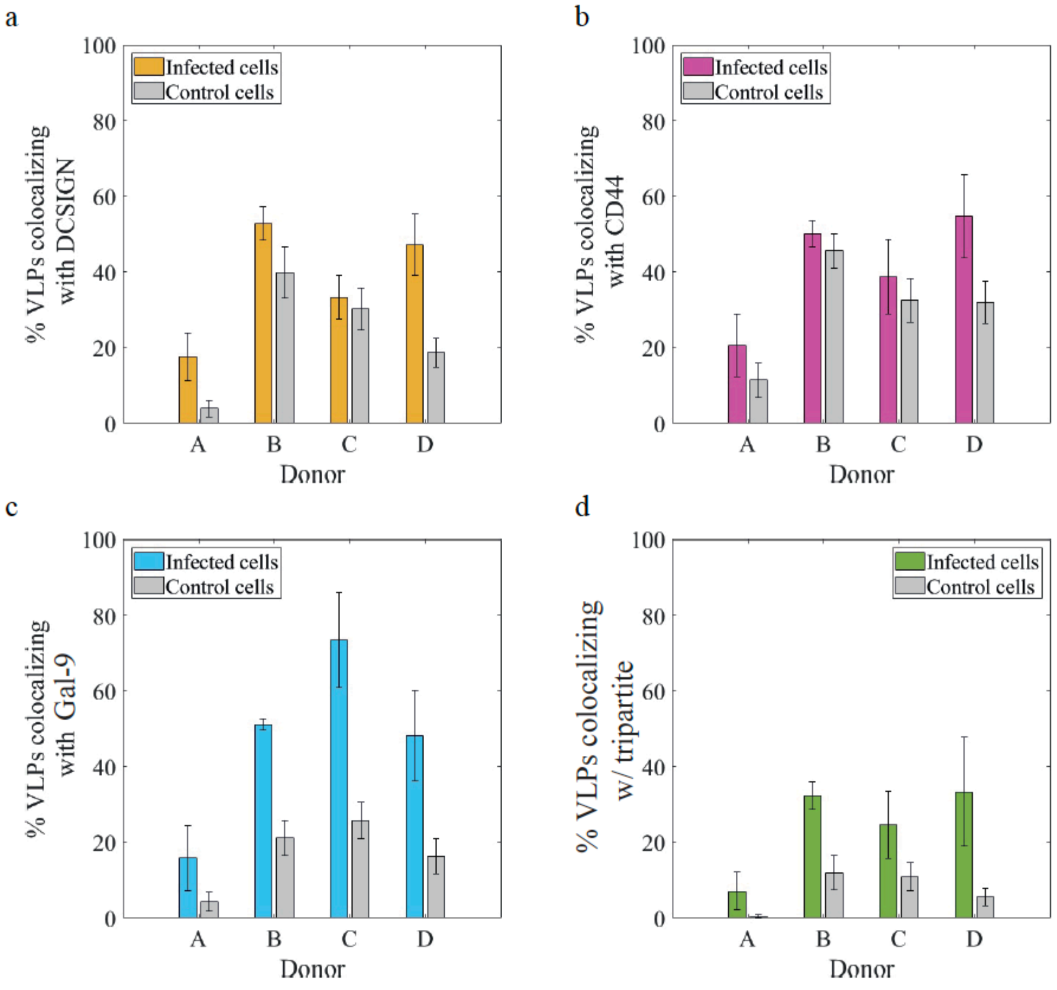


### SUPPLEMENTARY FIGURE 4

**Supplementary Fig. 4**: ***In-silico*** **simulations of stochastic viral landings on the cell membrane.** Percentage of HIV-1 VLPs colocalizing with DC-SIGN (**a**), CD44 (**b**), Gal-

9 (**c**) and with all three proteins (**d**) in infected cells (color bars) and in control cells (grey bars). In the case of control cells, we took the detected HIV-1 VLPs clusters of localizations from the experimental data and distributed them randomly on the reconstructed HiDenMaps from the cells non-exposed to HIV-1 VLPs.

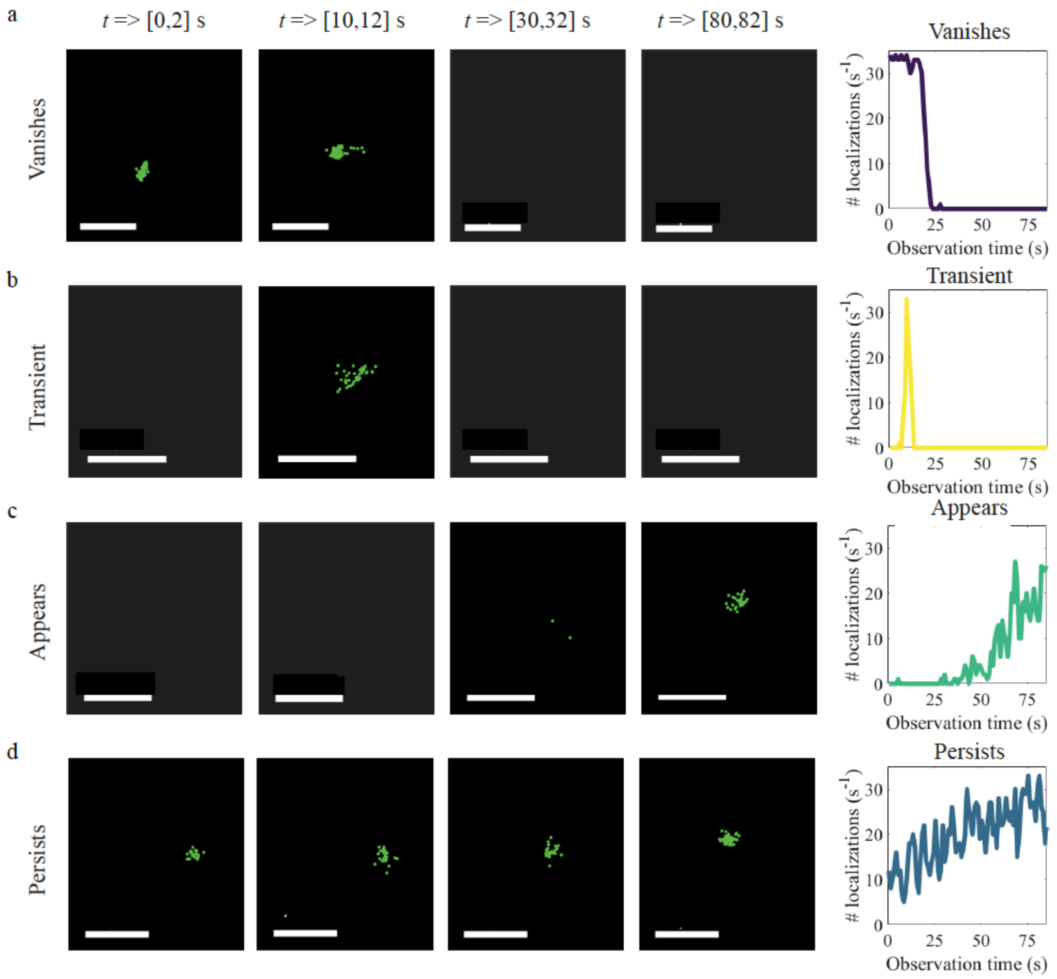


#### SUPPLEMENTARY FIGURE 5

**Supplementary Fig. 5: Classification of different VLP engagements on the cell membrane.** Representative HiDenMaps of VLP localizations generated at different time windows (2 seconds time-windows) showing distinctive viral engagements (see also the corresponding supplementary videos 7-10). The right column shows the number of VLP localizations of the corresponding HiDenMaps as a function of time. From top to bottom: **a**, The VLP *vanishes* from the field-of-view. **b,** The VLP arrives to the membrane but rapidly disengages and disappear, showing a *transient* engagement. **c,** The VLP *appears* during the observation time and remains engaged. **d,** The VLP shows *persistent* engagement to the cell membrane throughout the total observation time (90 seconds). Scale bars, 1 μm.

#### Captions for Supplementary videos:

**Supplementary Video 1:** Raw video of CD44 labeled with QD705 and imaged at a frame rate of 30 Hz for 90 s. The imaging is performed on live imDCs under TIR illumination. The scale bar is 10 μm.

**Supplementary Video 2:** Raw video of DC-SIGN labeled with QD655 and imaged at a frame rate of 30 Hz for 90 s. The imaging is performed on live imDCs under TIR illumination. The scale bar is 10 μm.

**Supplementary Video 3:** Raw video of Gal-9 labeled with QD605 and imaged at a frame rate of 30 Hz for 90 s. The imaging is performed on live imDCs under TIR illumination. The scale bar is 10 μm.

**Supplementary Video 4:** Overlay of the supplementary videos (1-3) keeping the source colors as follows: DC-SIGN (yellow), CD44 (magenta) and Gal-9 (cyan).

**Supplementary Video 5** Three color HiDenMap of DC-SIGN (yellow), CD44 (magenta) and Gal-9 (cyan). A time window of 2 seconds was used to cumulate the localizations of each protein.

**Supplementary Video 6:** Animation video of four-color single molecule trajectories, with DC-SIGN in yellow, CD44 in magenta, Gal-9 in cyan and, HIV-1 VLP in green. The frame rate for the imaging was 30 Hz and the acquisition was of 90 s. As the video shows, the VLP trajectory appears at later times where a pre-existing nanoplatform of the tripartite proteins was already found.

**Supplementary Video 7:** Four-color accumulated single-molecule localizations in time using a 2-seconds windows, showing the successful capture of a HIV-1 VLP by the tripartite proteins, DC-SIGN (yellow), CD44 (magenta) and Gal-9 (cyan). The localizations of each protein have been connected by the clustering algorithm. Scale bar: 1 μm. Note that this video shows a ROI of Supplementary Video 9.

**Supplementary Video 8:** Dynamic HiDenMap showing a VLP *vanishing* from the field-of-view due to unbinding or because the signal was too low to be detected. The time-window is set to 2 seconds, total observation time is 90 seconds. Scale bar is 1 μm. The color-coding is similar as to for Supplementary Video 4.

**Supplementary Video 9:** Dynamic HiDenMap showing a VLP with a *transient* behavior, i.e., the VLP arrives to the membrane but disengages and disappear. The time-window is set to 2 seconds, total observation time is 90 seconds. Scale bar is 1 μm. The color-coding is similar as to for Supplementary Video 4.

**Supplementary Video 10:** Dynamic HiDenMap showing a VLP *appearing* on the field-of-view, i.e., the VLP arrives at the plasma membrane and remains bound until the end of the acquisition time. The time window is set to 2 seconds, total observation time is 90 seconds. Scale bar is 1 μm. The color-coding is similar as to for Supplementary Video 4.

**Supplementary Video 11:** Dynamic HiDenMap showing a VLP with a *persistent* behavior, i.e., the VLP is visible throughout all the observation time (90 seconds). The time window is set to 2 seconds and the scale bar is 1 μm. The color-coding is similar as to for Supplementary Video 4.

**References**

1. Ester, M., Kriegel, H.-P., Sander, J. & Xu, X. A density-based algorithm for discovering clusters in large spatial databases with noise. kdd; 1996; 1996. p. 226-231.
